# Connecting Cancers to Chromosomal Locations Through Copy Number Variations

**DOI:** 10.1101/2023.10.03.560705

**Authors:** Saed Sayad, Mark Hiatt, Hazem Mustafa

## Abstract

**Background:** In 1960, scientists Peter Nowell and David Hungerford made a groundbreaking discovery by identifying the first genetic abnormality linked to cancer using cytogenetics. This discovery marked the beginning of research into many other genetic changes associated with cancer. These changes in chromosomes are like fingerprints, revealing how genes go haywire in cancer and destabilize the cell’s genetic makeup. In this paper, we focus on a clear connection between changes in the number of copies of certain genes at specific spots on chromosomes (called copy number variations or CNVs) and their role in various types of cancer.

**Method:** We obtained copy number data from various cancers cell lines through the DepMap portal website (https://depmap.org/portal/download/) and conducted diverse statistical analyses. Our approach involved comparing a single cancer cell line (primary disease) against the non-cancer cells. We used t-test to identify the top 100 genes that exhibited significant alterations in their copy number levels, both upregulated and downregulated. To enhance our understanding, we then linked these selected top 100 genes to their respective positions on the chromosomes.

**Results:** The top 100 upregulated genes associated with different cancers exhibit distinct chromosomal locations. For example all top 100 upregulated genes in Acute Myeloid Leukemia and Non-Hodgkin Lymphoma are on chromosome 17. Similarly, the top 100 downregulated genes linked to various cancers have their designated chromosomal locations.

**Conclusions:** Utilizing the DepMap copy number data, we have unveiled a distinctive linkage between cancer types and their specific chromosomal locales. The significance of these observed associations, when substantiated by biological evidence, holds the potential to significantly enhance our comprehension of cancer’s origins. Furthermore, such confirmation could pave the way for the development of more efficacious diagnostic, prognostic, and therapeutic strategies, promising a brighter outlook in the fight against cancer.

## Introduction

Cancer is a complex and genetically diverse disease, has been a subject of intense study since the pivotal year of 1960 when Peter Nowell and David Hungerford made a groundbreaking discovery in cytogenetics by identifying the first chromosomal abnormality associated with cancer (1). Subsequently, the field of cancer research has unraveled a multitude of chromosomal aberrations, numbering in the thousands, all intertwined with the disease (2). These aberrations encompass a spectrum of genetic changes, including insertions, deletions, inversions, substitutions, and duplications, which collectively serve as the hallmark of gene deregulation in cancer and culminate in genomic instability (3). Of particular significance is the role of copy number variations (CNVs) in the initiation and progression of numerous cancer types. In this study, we establish a definitive link between CNVs at specific gene locations on chromosomes and their widespread involvement in various cancer cell lines. This research sheds light on the intricate genetic underpinnings of cancer, with potential implications for diagnostics, prognostics, and therapeutic interventions.

## Data

We acquired the copy number data from the source: https://depmap.org/portal/download/. This dataset encompasses 1,740 unique cell lines and encompasses 27,562 individual genes. These cell lines originate from a diverse spectrum of 35 primary diseases, as depicted in Figure 1, which illustrates the distribution of each primary disease within the DepMap copy number dataset.

**Figure 1:**
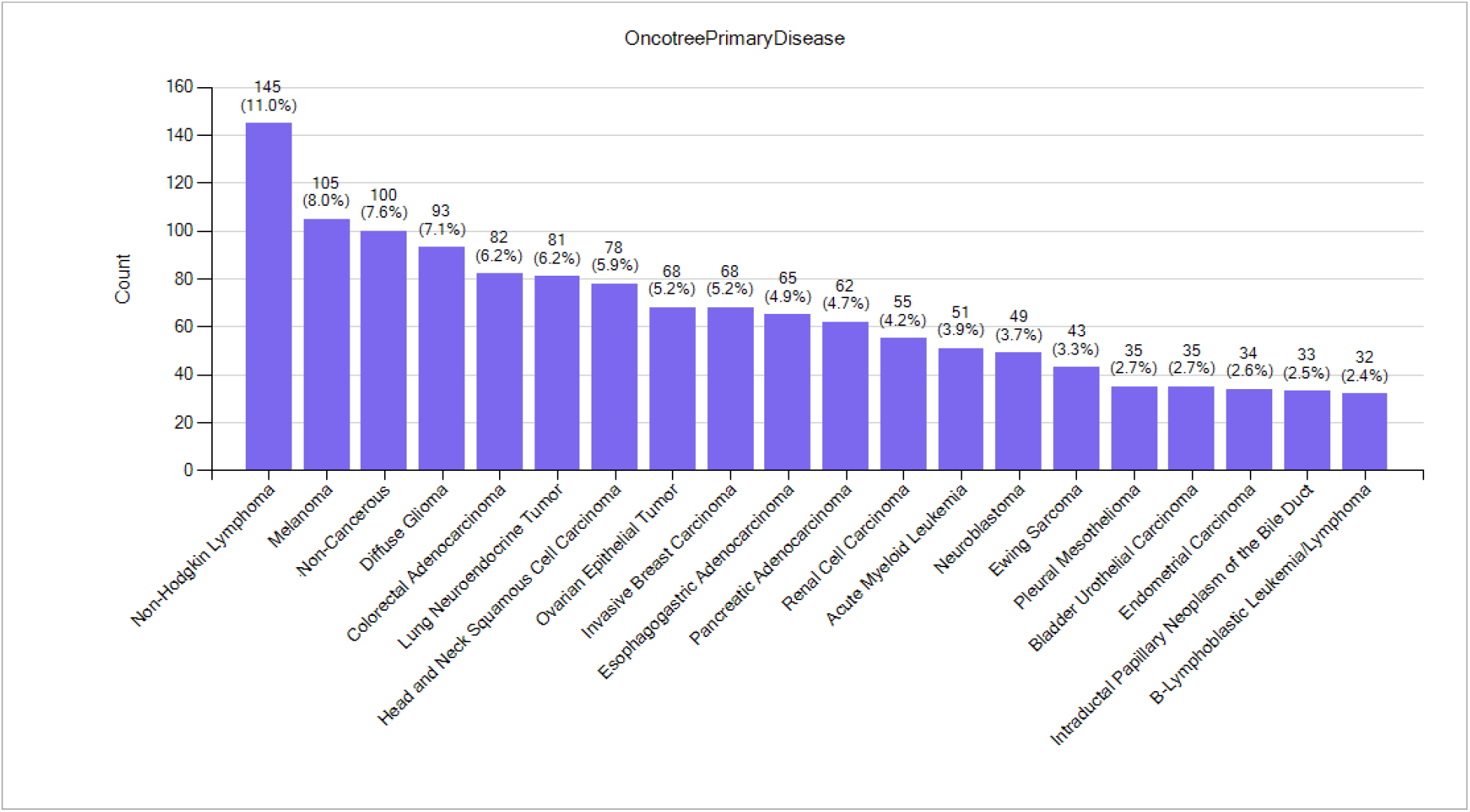
DepMap Primary Diseases (top 20 out of 82)

The table below presents a comprehensive list of the 82 primary diseases found within the DepMap Copy Number dataset.

## Data Analysis

We examined top 28 primary diseases cell lines against non-cancer cell lines using t-test to identify the 100 most significant genes that exhibited either upregulation or downregulation in each scenario. Subsequently, we determined the chromosomal locations of these selected 100 genes. Table 2 provides an overview of the primary chromosomal associations for the top 100 upregulated genes in each primary disease category (cancer cell lines). Notably, in instances such as Ewing Sarcoma and Osteosarcoma, all 100 upregulated genes were found to be situated on chromosome 1. Additionally, Table 3 highlights the predominant chromosome linked to the top 100 downregulated genes for each specific primary disease (cancer cell lines). For example, in cases like Acute Myeloid Leukemia and Non Hodgkin Lymphoma, all 100 downregulated genes were observed to reside on chromosome 17.

**Table 1:** 82 primary diseases found within the DepMap Copy Number dataset.

| **OncotreePrimaryDisease** | **Count** | **Count%** |
| --- | --- | --- |
| Non-Small Cell Lung Cancer | 152 | 8.39% |
| Non-Hodgkin Lymphoma | 145 | 8.00% |
| Melanoma | 105 | 5.79% |
| Non-Cancerous | 100 | 5.52% |
| Diffuse Glioma | 93 | 5.13% |
| Colorectal Adenocarcinoma | 82 | 4.53% |
| Lung Neuroendocrine Tumor | 81 | 4.47% |
| Head and Neck Squamous Cell Carcinoma | 78 | 4.30% |
| Invasive Breast Carcinoma | 68 | 3.75% |
| Ovarian Epithelial Tumor | 68 | 3.75% |
| Esophagogastric Adenocarcinoma | 65 | 3.59% |
| Pancreatic Adenocarcinoma | 62 | 3.42% |
| Renal Cell Carcinoma | 55 | 3.04% |
| Acute Myeloid Leukemia | 51 | 2.81% |
| Neuroblastoma | 49 | 2.70% |
| Ewing Sarcoma | 43 | 2.37% |
| Bladder Urothelial Carcinoma | 35 | 1.93% |
| Pleural Mesothelioma | 35 | 1.93% |
| Endometrial Carcinoma | 34 | 1.88% |
| Intraductal Papillary Neoplasm of the Bile Duct | 33 | 1.82% |
| B-Lymphoblastic Leukemia/Lymphoma | 32 | 1.77% |
| Esophageal Squamous Cell Carcinoma | 29 | 1.60% |
| Hepatocellular Carcinoma | 24 | 1.32% |
| Embryonal Tumor | 23 | 1.27% |
| T-Lymphoblastic Leukemia/Lymphoma | 22 | 1.21% |
| Rhabdomyosarcoma | 19 | 1.05% |
| Osteosarcoma | 18 | 0.99% |
| Myeloproliferative Neoplasms | 18 | 0.99% |
| Cervical Squamous Cell Carcinoma | 15 | 0.83% |
| Liposarcoma | 11 | 0.61% |
| Ocular Melanoma | 10 | 0.55% |
| Prostate Adenocarcinoma | 10 | 0.55% |
| Anaplastic Thyroid Cancer | 9 | 0.50% |
| Hodgkin Lymphoma | 9 | 0.50% |
| Chondrosarcoma | 8 | 0.44% |
| Cervical Adenocarcinoma | 7 | 0.39% |
| Well-Differentiated Thyroid Cancer | 7 | 0.39% |
| Intracholecystic Papillary Neoplasm | 7 | 0.39% |
| Synovial Sarcoma | 6 | 0.33% |
| Nerve Sheath Tumor | 6 | 0.33% |
| Cutaneous Squamous Cell Carcinoma | 6 | 0.33% |
| Breast Ductal Carcinoma In Situ | 5 | 0.28% |
| Rhabdoid Cancer | 5 | 0.28% |
| Ampullary Carcinoma | 4 | 0.22% |
| Uterine Sarcoma/Mesenchymal | 4 | 0.22% |
| Merkel Cell Carcinoma | 4 | 0.22% |
| Non-Seminomatous Germ Cell Tumor | 4 | 0.22% |
| Hepatoblastoma | 3 | 0.17% |
| Leiomyosarcoma | 3 | 0.17% |
| Fibrosarcoma | 3 | 0.17% |
| Meningothelial Tumor | 3 | 0.17% |
| Gestational Trophoblastic Disease | 3 | 0.17% |
| Undifferentiated Pleomorphic Sarcoma/Malignant Fibrous Histiocytoma/High-Grade Spindle Cell Sarcoma | 3 | 0.17% |
| Mucosal Melanoma of the Vulva/Vagina | 3 | 0.17% |
| Retinoblastoma | 2 | 0.11% |
| Urethral Cancer | 2 | 0.11% |
| Epithelioid Sarcoma | 2 | 0.11% |
| Adenosquamous Carcinoma of the Pancreas | 2 | 0.11% |
| Squamous Cell Carcinoma of the Vulva/Vagina | 2 | 0.11% |
| Small Bowel Cancer | 2 | 0.11% |
| Bladder Squamous Cell Carcinoma | 2 | 0.11% |
| Ovarian Cancer; Other | 1 | 0.06% |
| Head and Neck Carcinoma; Other | 1 | 0.06% |
| Ovarian Germ Cell Tumor | 1 | 0.06% |
| Breast Neoplasm; NOS | 1 | 0.06% |
| Myelodysplastic Syndromes | 1 | 0.06% |
| Prostate Small Cell Carcinoma | 1 | 0.06% |
| Hereditary Spherocytosis | 1 | 0.06% |
| Sarcoma; NOS | 1 | 0.06% |
| Poorly Differentiated Thyroid Cancer | 1 | 0.06% |
| Glassy Cell Carcinoma of the Cervix | 1 | 0.06% |
| Small Cell Carcinoma of the Cervix | 1 | 0.06% |
| Adrenocortical Carcinoma | 1 | 0.06% |
| Acute Leukemias of Ambiguous Lineage | 1 | 0.06% |
| Sex Cord Stromal Tumor | 1 | 0.06% |
| Chordoma | 1 | 0.06% |
| Medullary Thyroid Cancer | 1 | 0.06% |
| Pancreatic Neuroendocrine Tumor | 1 | 0.06% |
| Mixed Cervical Carcinoma | 1 | 0.06% |
| Salivary Carcinoma | 1 | 0.06% |
| Extra Gonadal Germ Cell Tumor | 1 | 0.06% |
| Hepatocellular Carcinoma plus Intrahepatic Cholangiocarcinoma | 1 | 0.06% |

**Table 2:** Chromosomal locations of the top 100 upregulated genes in 28 primary diseases.

| **Primary disease** | **Number of upregulated genes**<br>**(out of 100)** | **Chromosome** |
| --- | --- | --- |
| Ewing Sarcoma | 100 | 1 |
| Osteosarcoma | 100 | 1 |
| Cervical Squamous Cell Carcinoma | 100 | 5 |
| Hodgkin Lymphoma | 100 | 12 |
| Bile Duct Neoplasm | 100 | 17 |
| Neuroblastoma | 100 | 17 |
| T Lymphoblastic Leukemia | 100 | 17 |
| Bladder Urethelial Carcinoma | 100 | 20 |
| Head and Neck Squamous Cell Carcinoma | 100 | 20 |
| Esophagogastric Adenocarcinoma | 99 | 7 |
| Melanoma | 99 | 7 |
| Non Small Cell Lung Adenocarcinoma | 99 | 7 |
| Renal Cell Carcinoma | 99 | 7 |
| Ovarian Epithelial Tumor | 99 | 20 |
| Diffuse Glioma | 98 | 7 |
| Hepatocellular Carcinoma | 96 | 20 |
| B Lymphoblastic Leukemia | 95 | 1 |
| Endometrial Carcinoma | 94 | 1 |
| Well Differentiated Thyroid-Cancer | 94 | 17 |
| Myeloproliferative | 91 | 8 |
| Acute Myeloid Leukemia | 85 | 13 |
| Invasive Breast Cancer | 79 | 1 |
| Non Hodgkin Lymphoma | 76 | 1 |
| Pancreatic Adenocarcinoma | 75 | 20 |
| Colorectal Adenocarcinoma | 61 | 7 |
| Esophageal Squamous Cell Carcinoma | 60 | 5 |
| Prostate Adenocarcinoma | 53 | 1 |
| Lung Neuroendocrine Tumor | 44 | 3 |

**Table 3:** Chromosomal locations of the top 100 downregulated genes in 28 primary diseases.

| **Primary disease** | **Number of downregulated genes**<br>**(out of 100)** | **Chromosome** |
| --- | --- | --- |
| Head and Neck Squamous Cell Carcinoma | 100 | 3 |
| Lung Neuroendocrine Tumor | 100 | 3 |
| Bladder Urethelial Carcinoma | 100 | 4 |
| Hepatocellular Carcinoma | 100 | 4 |
| Acute Myeloid Leukemia | 100 | 17 |
| Non Hodgkin Lymphoma | 100 | 17 |
| Renal Cell Carcinoma | 99 | 14 |
| Melanoma | 98 | 10 |
| Neuroblastoma | 97 | 1 |
| Bile Duct Neoplasm | 97 | 4 |
| Pancreatic Adenocarcinoma | 97 | 18 |
| Non Small Cell Lung Adenocarcinoma | 96 | 8 |
| Ovarian Epithelial Tumor | 96 | 19 |
| Diffuse Glioma | 94 | 10 |
| Ewing Sarcoma | 94 | 14 |
| Invasive Breast Cancer | 92 | 8 |
| Well Differentiated Thyroid-Cancer | 88 | 2 |
| Esophageal Squamous Cell Carcinoma | 88 | 3 |
| Myeloproliferative | 79 | 9 |
| Colorectal Adenocarcinoma | 79 | 18 |
| Cervical Squamous Cell Carcinoma | 70 | 2 |
| Esophagogastric Adenocarcinoma | 68 | 19 |
| B Lymphoblastic Leukemia | 64 | 9 |
| Prostate Adenocarcinoma | 61 | 4 |
| Hodgkin Lymphoma | 61 | 13 |
| Endometrial Carcinoma | 56 | 4 |
| T Lymphoblastic Leukemia | 56 | 9 |
| Osteosarcoma | 27 | 19 |

## Discussion

By harnessing the wealth of information contained in the DepMap copy number data, we’ve been able to uncover an intersting connection between different types of cancer and the specific locations on chromosomes where these anomalies occur. This discovery is noteworthy because it suggests a potential link between genetic aberrations and the development of specific cancer types. The significance of these observed associations becomes even more profound when we can bolster them with solid biological evidence. When we can substantiate these findings through further research and experimentation, it has the potential to revolutionize our understanding of how cancer originates and progresses.

In practical terms, this could lead to several important advancements in the field of oncology. First and foremost, it could yield more accurate and effective diagnostic tools, enabling healthcare professionals to identify and classify cancer types with greater precision. This, in turn, would allow for more tailored treatment approaches, as we would have a deeper understanding of the underlying genetic factors driving the disease in each patient. Moreover, these findings might also enhance our ability to predict the course of the disease (prognosis) and develop novel therapeutic strategies. With a more comprehensive grasp of the genetic underpinnings of cancer, researchers and clinicians can potentially identify new targets for drug development and treatment interventions.

## Summary

The revelations stemming from the analysis of DepMap copy number data provide a pathway toward a brighter future where we can diagnose cancer more accurately, predict its behavior more effectively, and develop treatments that are tailored to individual patients, all of which could ultimately lead to improved outcomes and a higher quality of life for those affected by this devastating disease.

